# Utilization of Landscape of Kinases and Phosphosites To Predict Kinase-Substrate Association

**DOI:** 10.1101/2022.04.22.489231

**Authors:** Marzieh Ayati, Serhan Yılmaz, Filipa Blasco Tavares Pereira Lopes, Mark R. Chance, Mehmet Koyutürk

## Abstract

**Motivation:** Protein phosphorylation is a key post-translational modification that plays a central role in many cellular processes. With recent advances in biotechnology, thousands of phosphorylated sites can be identified and quantified in a given sample, enabling proteome-wide screening of cellular signaling. However, the kinase(s) that phosphorylate most (> 90%) of the identified phosphorylation sites are unknown. Knowledge of kinase-substrate associations is also mostly limited to a small number of well-studied kinases, with 20% of known kinases accounting for the phosphorylation of 87% of currently annotated sites. The scarcity of available annotations calls for the development of computational algorithms for more comprehensive and reliable prediction of kinase-substrate associations.

**Results:** To broadly utilize available structural, functional, evolutionary, and contextual information in predicting kinase-substrate associations, we develop a network-based machine learning framework. Our framework integrates a multitude of data sources to characterize the landscape of functional relationships and associations among phosphosites and kinases. To construct a phosphosite-phosphosite association network, we use sequence similarity, shared biological pathways, co-evolution, co-occurrence, and co-phosphorylation of phosphosites across different biological states. To construct a kinase-kinase association network, we integrate protein-protein interactions, shared biological pathways, and membership in common kinase families. We use node embeddings computed from these heterogeneous networks to train machine learning models for predicting kinase-substrate associations. Our systematic computational experiments using the PhosphositePLUS database shows that the resulting algorithm, NetKSA, outperforms state-of-the-art algorithms and resources, including KinomeXplorer and LinkPhinder, in reliably predicting KSAs. By stratifying the ranking of kinases, NetKSA also enables annotation of phosphosites that are targeted by relatively less-studied kinases. Finally, we observe that the performance of NetKSA is robust to the choice of network embedding algorithms, while each type of network contributes valuable information that is complementary to the information provided by other networks.

**Conclusion:** Representation of available functional information on kinases and phosphorylation sites, along with integrative machine learning algorithms, has the potential to significantly enhance our knowledge on kinase-substrate associations.

**Availability:** The code and data are available at compbio.case.edu/NetKSA.

## 1 Introduction

Protein phosphorylation is one of the most important post-translational modifications that play an important role in cellular signaling. Phosphorylation involves phospho-proteins whose activity can be altered by the phosphorylation of their specific sites (a.k.a substrate), kinases that phosphorylate the phosphoproteins at specific sites, and phosphatases that dephosphorylate these proteins. Dysregulation of the kinase-substrate associations are regularly observed in complex diseases, including cancer. Therefore, kinases have emerged as an important class of drug targets for many diseases [1].

Recent advances in mass spectrometry (MS) based technologies drastically enhanced the accuracy and coverage of phosphosite identification and quantification. However, most identified phosphosites do not have kinase annotations, and large scale and reliable prediction of which kinase can phosphorylate which phosphosites remains challenging. In the last decade, several computational methods are developed to predict kinase-substrate associations (KSAs). The earlier KSA prediction methods focus mainly on sequence motifs recognized by the active sites of kinases.[2, 3, 4]. Later methods integrate other contextual information such as protein structure and physical interactions to improve the accuracy of prediction methods [5, 6]. Recently, we developed CophosK [7], the first kinase-substrate prediction algorithm that utilizes large-scale mass spectrometry based phospho-proteomic data to incorporate contextual information. While these tools improve the kinase-substrate associations prediction, the knowledge about the substrates of kinases is still unequally distributed, where 87% of phosphosites are assigned to 20% of well-studied kinases [8].

In parallel, machine learning algorithms that utilize network models gain significant traction in computational biology [9, 10]. Inspired by these developments, we here develop a comprehensive framework for integrating broad functional information on kinases and phospho-proteins to build machine learning models for predicting kinase-substrate associations. Our framework uses heterogeneous network models to represent the functional relationships between phosphorylation sites, as well as kinases. Namely, we integrate structural, evolutionary, functional, and contextual information to characterize the landscape of functional relationships and associations among phosphosites and kinases. Since MS-based phosphoproteomic data can present a relatively unbiased view of signaling states, we also incorporate co-occurrence and co-phosphorylation across multiple MS-based phospho-proteomic studies into network construction. After constructing phosphosite association and kinase association networks, we use node embedding algorithms to derive low-dimensional vector representations for phosphosites and kinases, which are in turn used to train machine learning models.

We systematically investigate the predictive performance of reliability of the proposed framework, NetKSA, using established kinase-substrate associations from PhosphositePLUS. Using a cross-validaton framework in two problem settings (link prediction and prioritization), we investigate the effect of the network embedding algorithms, the contribution of different types of networks, the value added by network topology, and compare the performance of NetKSA against state-of-the-art algorithms. In order to mitigate the bias toward well-studied kinases in the KSA prediction [11], we propose a kinase stratification strategy based on the number of known substrates. Our results show that NetKSA, outperforms state-of-the-art methods in overall prediction performance. Finally, we observe that the performance of NetKSA is robust to the choice of network embedding algorithms, while each type of network contributes valuable information that is complementary to the information provided by other networks.

## 2 Materials and Methods

The workflow of the proposed framework for kinase-substrate association prediction is shown in Figure 1. As seen in the figure, we first construct two networks, one to model the functional relationship between phosphorylation sites. For this purpose, we incorporate available knowledge on functional associations between phosphosites, and between kinases, and integrate these knowledge into creating two networks of phosphoSite Association Network (SAN) and Kinase Association Network (KAN). Subsequently, we apply node embedding method to reduce the number of dimension and present the nodes of networks with lower dimensions. We use the known kinase-substrate association obtained from PhosphoSitePLUS to train the predictive models to predict kinase-substrate associations.

**Figure 1:**
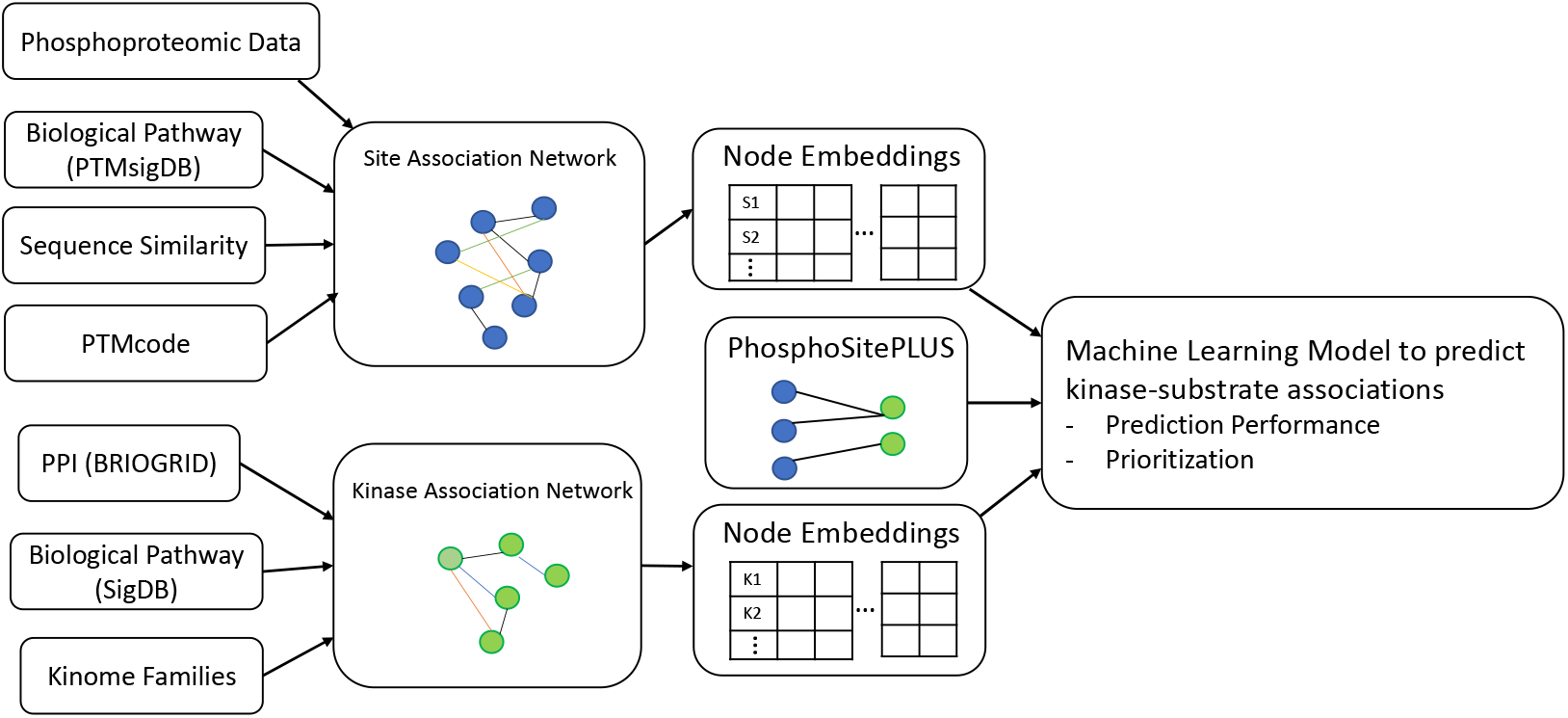
Workflow of NetKSA. We first construct two networks to represent the functional relationships and associations among phosphosites and kinases. The nodes in site association network represent phosphosites, and the edges represents (1) co-evolution obtained from PTMcode, (2) sequence similarity obtained from the sequence alignment, (3) shared biological pathways, (4) co-occurrence across 86 studies published in PhosphositePLUS, and (5) co-phosphorylation across samples in 9 MS-based phosphoprotoemics datasets. The nodes in the kinase association network represent kinases and the edges represent (1) physical interactions between kinases, (2) shared biological pathways, and (3) kinases that are in the same family. After construction of networks, we use node embedding algorithms on each network to compute a low-dimensional representation for each node. We then use the kinase-substrate associations (KSAs) obtained from PhosphoSitePLUS to train machine learning models for predicting KSAs.

### 2.1 PhosphoSite Association Network (SAN)

We define a phosphoSite Association Network (SAN) as a network that represents *potential* functional relationships between pairs of phosphosites. In SAN network *G_s_*(*V,E*), *V* denotes the set of nodes in the network, each of which represents a phosphosite. The edge set *E* denotes the set of pairwise functional relationships between phosphosites, where an edge *s_i_s_j_* ∈ *E* between phosphosites *s_i_, s_j_* ∈ *V* may represent one of the following relationships:

- **Functional, Evolutionary, and Structural Association between Phosphosites.** PTMCode is a database of known and predicted functional associations between phosphorylation and other post-translational modification sites [12]. The associations included in PTMCode are curated from the literature, inferred from residue co-evolution, or are based on the structural distances between phosphosites. We utilize PTMcode as a direct source of functional, evolutionary, and structural associations between phosphorylation sites.
- **Sequence Similarity**. We downloaded the sequences within ±7 residues around each site in the protein sequence from phosphoSitePLUS, and perform sequence alignment using BLOSUM62 scoring method. There is an edge between two sites *s_i_* and *s_j_* if their distance is less than 3 standard deviation around the average across all pairs of sites.
- **Biological Pathways**. We use PTMsigDB as a reference database of site-specific phosphorylation signatures of kinases, perturbations, and signaling pathways [13]. While PTMSigDB provides data on all post-translational modifications, we here use the subset that corresponds to phosphorylation. There area 2398 phosphosites that are associated with 388 different perturbation and signaling pathways. We represent these associations as a binary network of signaling-pathway associations among phosphosites, in which an edge between two phosphosites indicates that the phosphorylation of the two sites is involved in the same pathway.
- **Co-Occurrence**. Li et al showed that phosphorylated sites that are modified together tend to participate in similar biological process [14]. We use the high-throughput MS analyses across 88 different conditions from phosphoSitePLUS [15] which includes 16 human tissue as well as 28 cultural cell lines and 44 disease cells. We put an edge between two sites *s_i_* and *s_j_* if the p-value of their co-occurrence is less than 0.005.
- **Co-Phosphorylation**. Since co-phosphorylation can potentially capture context-specific, as well as universal functional relationships among phosphorylation sites, we also incorporate the high co-phosphorylated sites in the edges of the network. For this purpose, we use 9 MS-based phosphorptoemics data with enough number of dimensions (i.e. states) in order to obtain a reliable co-phosphorylation [7]. We calculated the integrated co-phosphorylation between each pair of sites as following:

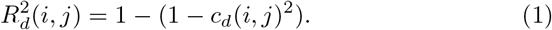 Here, *c_d_*(*i,j*) denotes the co-phosphorylation (measured by Biweight-midcorrealtion) in dataset *d* ∈ *D*. There is an edge between two sites *s_i_* and *s_j_* if the absolute value of their co-phosphorylation is larger than 2 standard deviation of the average across all pairs of sites. In our experiment, the MS-based phosphoproteomics data includes three breast cancer data [16, 17, 18], two ovarian cancer data [19, 17], one colorectal cancer [20], one lung cancer [21], one Alzheimer’s disease [22] and one retinal pigmented eputhelium data [23].

### 2.2 Kinase Association Network (KAN)

We define a Kinase Association Network (KAN) as a network that represents functional relashiptnshipd between pairs of kinases. In KAN network *G_k_* (*V, E*), *V* denotes the set of nodes in the network, each of which represents a kinase. The edge set *E* denotes the set of pairwise functional relationships between kinases, where an edge *k_l_k_r_* ∈ *E* between kinases *k_l_,k_r_* ∈ *V* may represent one of the following relationships:

- **Protein-Protein Interaction (PPI)**. If two kinases *k*_ℓ_ and *k_r_* physically interact, then there is an edge between *k*_ℓ_ and *k_r_* in the KAN. In our experiments, we use the PPIs that are annotated as ”physical” in the BIOGRID PPI database [24] to infer the PPI edges in the network.
- **Biological Pathways**. If two kinases *k*_ℓ_ and *k_r_* are reported to be in the same pathway, then there is an edge between *k*_ℓ_ and *k_r_* in the KAN. In our experiments, we use mSigDB which is the collection of canonical pathways and experimental signatures as a reference [25].
- **Kinome Families**. If two kinases *k*_ℓ_ and *k_r_* belongs to the same family according to the human kinome [26], then there is an edge between them.

### 2.3 Computing Network Profile for Sites and Kinases

To calculate the network profile of a given node *v* ∈ *V* in the given network *G* = (*V, E*), we utilize the node embedding approach. Given a graph *G*, a graph embedding is a mapping *f* : *v_i_* → *y_i_* ∈ *R^d^* such that *d* ≪ |*V*| and the function *f* preserves some proximity measure defined on graph *G*. In other words, a node embedding map each node to a low-dimensional feature vector and tries to preserve the connection strength between nodes. In our experiment, we use different node embedding methods including node2vec [27], DNGR [28], Deepwalk [29], and Verse [30]. These methods use different approach for the embedding. For example, node2vec and DeepWalk use random walk, Verse uses neural network and DNGR uses deep learning method. We calculate the node embedding as network profiles for every site *s_i_* in *G_s_* and every kinase *k*_l_ in *G_k_* using the default parameters in each method.

### 2.4 Predicting Kinase-Substrate Associations

We use the sets of known KSAs obtained from PhosphoSitePLUS (PSP) [15] as a positive reference. We generate the negative training set using random selection of the pair of sites and kinases that are not reported to be associated in PSP. To train the models, we use the network profiles of site-kinase pairs by using concatenation approach: 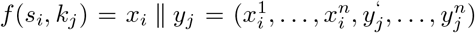 where *x_i_* ∈ *R^n^* and *y_j_* ∈ *R^n^* are the network profile of *s_i_* and *k_j_* in their associated network, respectively. We tackle the KSA prediction with two approach:

I. **Link Prediction**. In this approach, we formulate the problem as binary classification and link prediction between kinases and sites: Given two networks *G_s_* and *G_k_*, as well as a pair of nodes (*s_i_, k_l_*), our objective is to evaluate the existence of the link between *s_i_* and *k_l_*. We use Random Forest, and 5-fold cross validation in this analysis. We assess the overall performance of the method using area of the ROC curve (AUC).
II. **Prioritization**. In this approach, we formulate the problem as kinase prioritization: Given two networks *G_s_* and *G_k_*, and a site *s_i_*, our objective is to compute an association score between *s_i_* and all the kinases, and rank the kinases based on their association score. We use Random Forest and leave-one-out validation in this analysis. We assess the performance of the method using hit@k accuracy. This metric evaluated the performance in terms of the number of times in which the actual kinases of sites are ranked in the top *k*, where *k* ∈ {1, 5, 10, 20}.

### 2.5 Stratifying Kinases to Improve the Prediction Performance

In order to study the bias in the KSA predictions toward the more well-studies kinases. We categorize the kinases in to three categories: (i) The poor kinases where their number of known associated substrates is less than 5, (ii) The average kinases, where their number of known associated substrates between 5 and 20, and (iii) The rich kinases where their number of known associated substrates is larger than 20. We evaluate the performance of the methods for overall and also each category of kinases, separately. We also keep the proportion of the category the kinases while generating the negative training set. In order to remove the bias from the predictive models, for the prioritization task, we limit the ranking to the kinases in each category. The premise of this approach is that the kinases in each category should just compete with the same category kinases to have a better chance to rank high for their associated substrate.

## 3 Results and Discussion

We use PhosphoSitePLUS as a reference dataset for kinase-substrate associations (KSAs) [15]. Considering the phosphosites and kinases in our networks, we use 2083 KSAs from PhosphositePLUS in our computational experiments. To evaluate the performance of the kinase-substrate association prediction method, we limit the site network to the known substrates obtained from PhosphoSitePLUS. We remove the individual nodes that are not connected to any other nodes from both of the networks. The number of sites and edges in the final kinase-kinase and phosphosite-phosphosite association networks and their types are shown in Figure **??**(a). The overlaps between different types of association networks are shown in Figure **??**(b). The low overlap between different phosphosite-phosphosite association networks suggests that all different types of networks provide information that are potentially complementary with each other.

### 3.1 Kinase-Substrate Association as Link Prediction

We first use different embedding methods, and 5-fold cross validation to evaluate the performance of NetKSA in predicting KSAs formulated as link prediction. In our computational experiments, we consider different numbers of embedding dimensions (Figure S1) and find out that *d* = 16 is optimal for all algorithms considered, thus we perform all remaining experiments using 16 dimensions for the embedding vectors.

The link prediction performance of NetKSA using different embedding algorithms is presented in Figure 2(a). We evaluate the performance for all the KSAs, as well as KSAs that its kinase belongs to different category (i.e. poor, average, rich) separately. In this analysis, there are 103 kinases in the poor category (*δ* < 5), 64 kinases in the average category (5 ≤ *δ* < 20), and 21 kinases in the rich category (*δ* ≥ 20) (the rest of kinases in the kinase-kinase association network do not have any target sites that are present in the site-site association network). These kinases corresponds to 218 KSAs in poor category, 613 KSAs in the average category and 1252 KSAs in the rich category. The negative set for the training of the model is randomly generated while keeping the proportion of KSA categories. The bar plots show the average across 10 runs. As seen in the figure, the prediction performance highly depends on the the kinase category and the AUC observed by considering all kinases together closely follows the prediction performance for rich kinases. This observation demonstrates the importance of performing stratified analyses to accurately characterize the performance of KSA prediction as a function of what is already known about the kinase and characterize the bias in algorithms.

**Figure 2:**
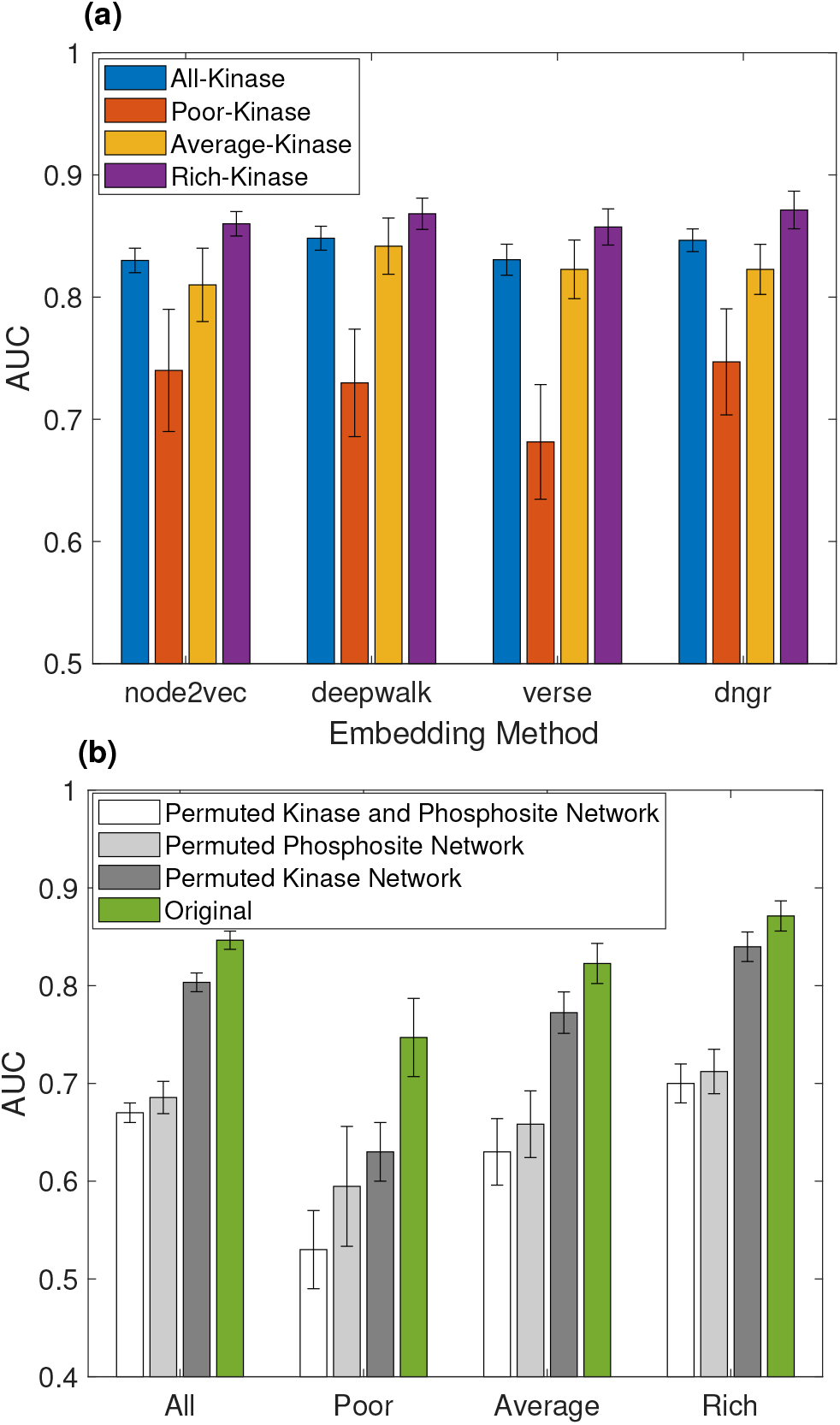
The contribution of embedding algorithms and functional networks on KSA prediction performance. (a) The AUC of the predictions of NetKSA using four different node embedding algorithms. For each embedding algorithm, the AUC is shown for all KSAs (blue bar), the KSAs where the kinase belongs to the poor category (red), the average category (gold), and rich category (purple). (b) The prediction performance of NetKSA using DNGR for node embedding using real vs. randomized networks. AUC on the real kinase-kinase and phosphosite-phosphosite association networks (green bar), when only the kinase association network is randomly permuted by preserving node degrees (dark grey), when only the site association network is permuted by preserving node degrees (light grey), when both networks are permuted (white). Each bar shows the average AUC across 10 runs and the error bar shows standard deviation.

As seen in Figure 2(a), the prediction performance of NetKSA is robust to the choice of network embedding algorithms. We select DNGR for further analyses due to its slightly better overall performance that is also most balanced across different kinase categories.

To evaluate the value added by the network to the prediction, we randomly permute site association and kinase association networks while preserving the degree distribution and apply NetKSA by using the permuted networks in place of the actual networks. The results of this analysis are presented in Figure 2(b). As seen in the figure, the prediction performance using original networks is one or more standard deviation(d) above the prediction performance of the method when using permuted networks. This result shows the networks contribute valuable information for KSA prediction. Importantly, randomization of the prediction performance declines more when the phosphiste network is permuted, suggesting that the functional information on the phosphosites provides significant and specific information on the kinase(s) that target(s) the phosphosites.

It is also interesting that the poor kinase category benefits the most in comparison with other categories when the original networks are used. This shows that the information provided by functional associations among sites and kinases reduce the gap between under-studied and well-studied kinases. Note that the models that are based on permuted networks perform better than what would be expected at random, suggesting that these models can learn bias in the benchmarking data to appear as if they are learning what they are designed to learn. However, the performance of the model that is trained on both permuted networks is equal to what would be expected at random for poor kinases, demonstrating that the validation strategy we employ here (stratification of kinases and comparison against permuted networks) provides significant insights into what these models actually learn.

### 3.2 Contribution of Different Networks on Prediction Performance

In order to evaluate the contribution of different types of networks in capturing the landscape of functional association among phosphosites and kinases, we evaluate the performance of KSA predictions using different networks. For this analysis, we perform KSA prediction using 5-fold cross validation, by adding one network at a time to the integrated network of kinase-kinase and phosphosite-phosphosite associations, while keeping the other network fully integrated. The results of this analysis are shown in Figure 3. As seen in the figure, as we add different types of functional information for the sites and kinases, the prediction performance improves. Importantly, the new networks add information about the the individual sites and kinases and connect them to other nodes, and consequently increase the coverage of kinases, sites and kinase-substrate associations for which predictions can be made. Finally, we observe that the information contributed by different phosphosite networks is more complementary to each other as compared to the kinase networks, which is not surprising as the overlap between these networks is also considerably low.

**Figure 3:**
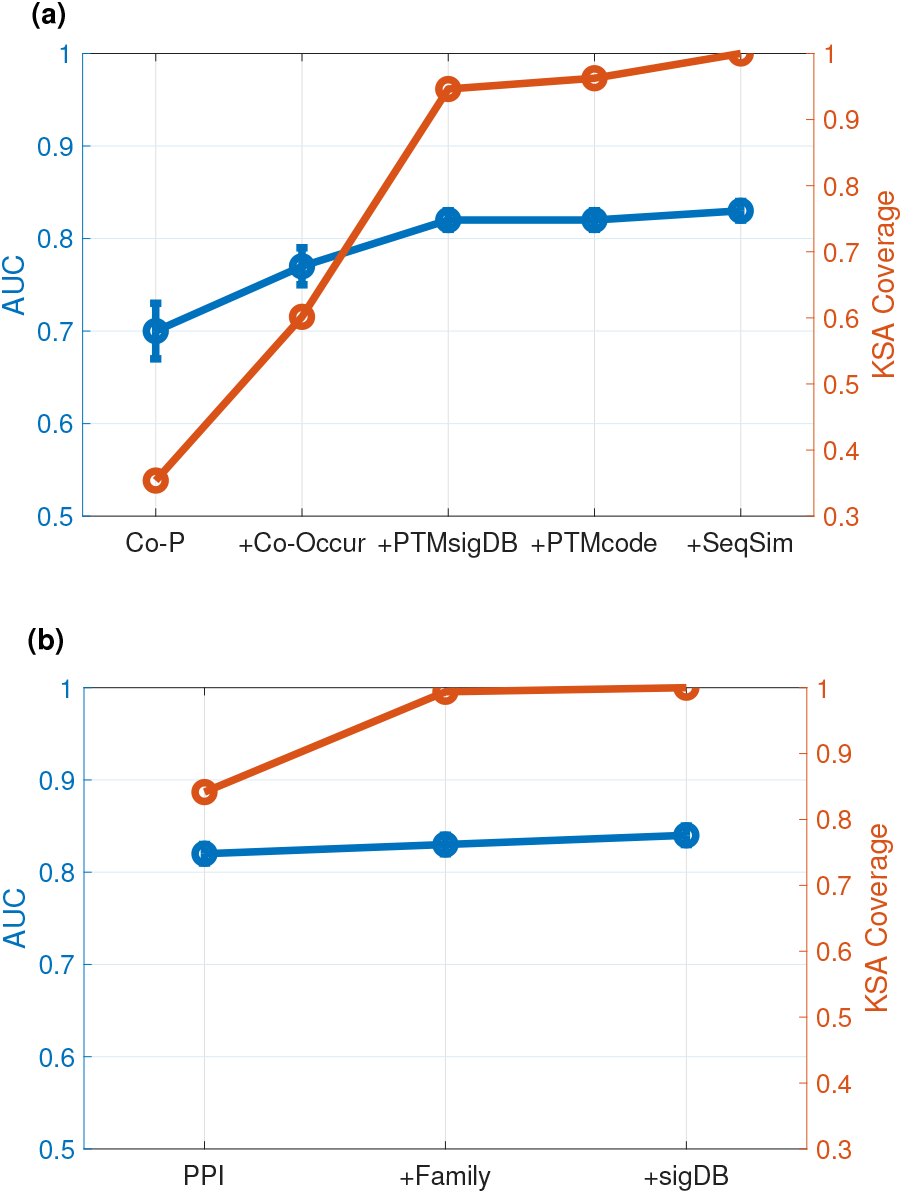
Contribution of different types of networks on the prediction of KSAs. The cumulative effect of each (a) phosphosite-phosphosite association network and (b) kinase-kinase association network on the AUC of predictions (left y axis; blue), and the coverage of kinase-substrate associations (right y axis; red) - the fraction of KSAs for which both the kinase and the site are present in the integrated network so that a prediction can be made.

### 3.3 Prioritization of Kinases for Phosphorylation Sites

To test the effectiveness of our method, we use leave-one-out cross validation. Namely, for each phosphosite, we hide the association between phosphosite and its known kinase (called the target kinase), and we use other reported KSAs to rank the likely kinases for that phosphosite. For this analysis, we use dngr as the embedding method and random forest with 100 classification trees as the score prediction model. For each phosphosite, we rank all kinases based on the calculated score and determine the rank of the target kinase across all kinases. If the target kinase is within the top *k* ∈ {1, 5, 10, 20}, it is considered a the true positive.

We compare our method with two other state-of-the-art methods, KinomeXplorer and LinkPhinder, that also use the network for KSA prediction. KinomeXplorer [5] utilizes the sequences match scoring and network proximity of kinases and substrates to predict KSAs. It is an improved version of NetworKIN[4] and NetPhorest [31]. LinkPhinder [32] is also another predictive model that utilizes the motif characteristics to create a knowledge graph and uses statistical relational learning and node embedding to predict KSAs. The result of this analysis is presented in Figure 4. As seen in the figure, the proposed method with kinase stratification outperform all methods in overall prediction performance, and also average and rich categories. For the poor kinases, the LinkPhinder presents a better result for top 1 and top 5 ranking. Both proposed method and LinkPhinder present a better performance compare to KinomeXplorer for poor kinases.

**Figure 4:**
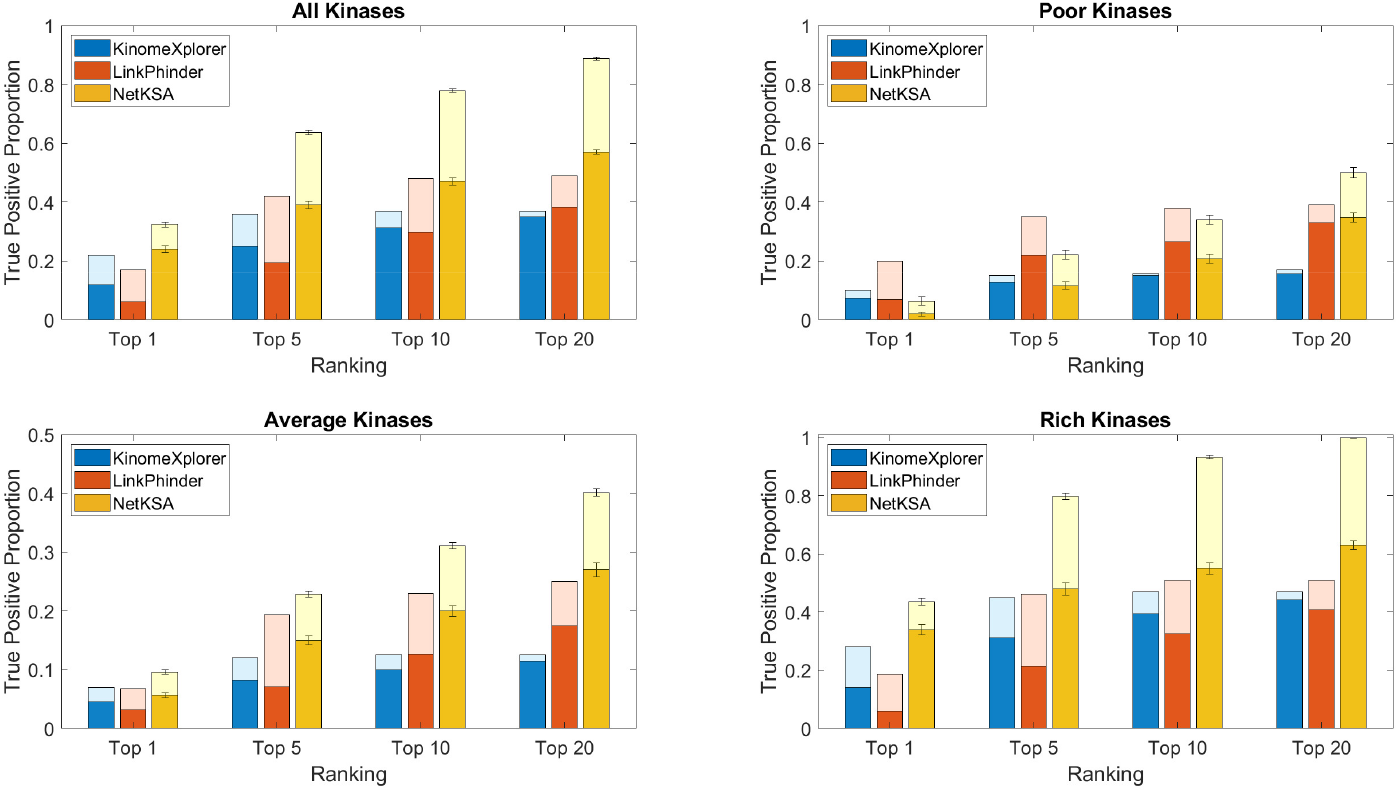
Performance of NetKSA, KinomeXplorer and LinkPhinder in prioritizing kinases for a given phosphosite. For each phosphosite, we perform leave-one-out cross validation by hiding the association between the phosphosite and one of its associated kinases (target kinase) to rank the likely kinases for the phosphosite using KinomeXplorer(blue), LinkPhinder(red), and proposed method using constructed networks (gold). We report the fraction of phosphosites for which the target kinase is ranked in the top 1, top 5, top 10 and top 20 predicted kinases by each method. For each bar, the dark section presents the result when all the kinases are ranked together, and the light section presents the improvement of performance when the target kinase is ranked within its category (with stratification). Each panel presents the performance on each category of kinases: poor (*δ* < 5), average(5 ≤ **δ** < 20), and rich (*δ* ≥ 20) kinases (as indicated in each panel).

#### 3.3.1 Kinase Stratification

In the kinase prioritization, we rank the kinases in each category (i.e poor, average, rich) separately, and determine if the target kinase is ranked in top *k* of its category. The premise of this approach is that the kinase that are understudied does not to compete with the well-studies kinases. Using kinase stratification, the hypothesis is that it is more likely that the target kinase wins the competition in ranking compare to the kinases in its own category. We apply this strategy on NetKSA and also KinomeXplorer and LinkPhinder. The result of this analysis is presented in Figure 4. For each bar in the figure, the dark section is the performance without kinase stratification, and the light-color section is the improvement of the performance using the kinase stratification.

## References

[1] Fleur M Ferguson and Nathanael S Gray. Kinase inhibitors: the road ahead. Nature reviews Drug discovery, 17(5):353–377, 2018.

[2] Pawel Durek, Christian Schudoma, Wolfram Weckwerth, Joachim Selbig, and Dirk Walther. Detection and characterization of 3d-signature phosphorylation site motifs and their contribution towards improved phosphorylation site prediction in proteins. BMC bioinformatics, 10(1):1–17, 2009.

[3] John C Obenauer, Lewis C Cantley, and Michael B Yaffe. Scansite 2.0: Proteome-wide prediction of cell signaling interactions using short sequence motifs. Nucleic acids research, 31(13):3635–3641, 2003.

[4] Rune Linding, Lars Juhl Jensen, Gerard J Ostheimer, Marcel ATM van Vugt, Claus Jørgensen, Ioana M Miron, Francesca Diella, Karen Colwill, Lorne Taylor, Kelly Elder, et al. Systematic discovery of in vivo phosphorylation networks. Cell, 129(7):1415–1426, 2007.

[5] Heiko Horn, Erwin M Schoof, Jinho Kim, Xavier Robin, Martin L Miller, Francesca Diella, Anita Palma, Gianni Cesareni, Lars Juhl Jensen, and Rune Linding. Kinomexplorer: an integrated platform for kinome biology studies. Nature methods, 11(6):603–604, 2014.

[6] Majbrit Hjerrild, Allan Stensballe, Thomas E Rasmussen, Christine B Kofoed, Nikolaj Blom, Thomas Sicheritz-Ponten, Martin R Larsen, Søren Brunak, Ole N Jensen, and Steen Gammeltoft. Identification of phosphorylation sites in protein kinase a substrates using artificial neural networks and mass spectrometry. Journal of proteome research, 3(3):426–433, 2004.

[7] M. Ayati, D Wiredja, D Schlatzer, S. Maxwell, Ming Li, M. Koyuturk, and M.R Chance. Cophosk: A method for comprehensive kinase substrate annotation using co-phosphorylation analysis. PLOS computational biology, 2019.

[8] Elise J Needham, Benjamin L Parker, Timur Burykin, David E James, and Sean J Humphrey. Illuminating the dark phosphoproteome. Science signaling, 12(565):eaau8645, 2019.

[9] Thomas Gaudelet, Ben Day, Arian R Jamasb, Jyothish Soman, Cristian Regep, Gertrude Liu, Jeremy BR Hayter, Richard Vickers, Charles Roberts, Jian Tang, et al. Utilizing graph machine learning within drug discovery and development. Briefings in bioinformatics, 22(6):bbab159, 2021.

[10] Giulia Muzio, Leslie O’Bray, and Karsten Borgwardt. Biological network analysis with deep learning. Briefings in bioinformatics, 22(2):1515–1530, 2021.

[11] Iman Deznabi, Busra Arabaci, Mehmet Koyuturk, and Oznur Tastan. Deepkinzero: zero-shot learning for predicting kinase-phosphosite associations involving understudied kinases. Bioinformatics, 36(12):3652–3661, 2020.

[12] Pablo Minguez, Ivica Letunic, Luca Parca, Luz Garcia-Alonso, Joaquin Dopazo, Jaime Huerta-Cepas, and Peer Bork. Ptmcode v2: a resource for functional associations of post-translational modifications within and between proteins. Nucleic acids research, 43(D1):D494–D502, 2014.

[13] Karsten Krug, Philipp Mertins, Bin Zhang, Peter Hornbeck, Rajesh Raju, Rushdy Ahmad, Matthew Szucs, Filip Mundt, Dominique Forestier, Judit Jane-Valbuena, et al. A curated resource for phosphosite-specific signature analysis. Molecular & cellular proteomics, 18(3):576–593, 2019.

[14] Ying Li, Xueya Zhou, Zichao Zhai, and Tingting Li. Co-occurring protein phosphorylation are functionally associated. PLoS computational biology, 13(5):e1005502, 2017.

[15] Peter V Hornbeck, Bin Zhang, Beth Murray, Jon M Kornhauser, Vaughan Latham, and Elzbieta Skrzypek. Phosphositeplus, 2014: mutations, ptms and recalibrations. Nucleic acids research, 43(D1):D512–D520, 2014.

[16] Kuan-lin Huang, Shunqiang Li, Philipp Mertins, Song Cao, Harsha P Gunawardena, Kelly V Ruggles, DR Mani, Karl R Clauser, Maki Tanioka, Jerry Usary, et al. Proteogenomic integration reveals therapeutic targets in breast cancer xenografts. Nature communications, 8:14864, 2017.

[17] Philipp Mertins, Feng Yang, Tao Liu, DR Mani, Vladislav A Petyuk, Michael A Gillette, Karl R Clauser, Jana W Qiao, Marina A Gritsenko, Ronald J Moore, et al. Ischemia in tumors induces early and sustained phosphorylation changes in stress kinase pathways but does not affect global protein levels. Molecular & cellular proteomics, 2014.

[18] Philipp Mertins, DR Mani, Kelly V Ruggles, Michael A Gillette, Karl R Clauser, Pei Wang, Xianlong Wang, Jana W Qiao, Song Cao, Francesca Petralia, et al. Proteogenomics connects somatic mutations to signalling in breast cancer. Nature, 534(7605):55, 2016.

[19] Hui Zhang, Tao Liu, Zhen Zhang, Samuel H Payne, Bai Zhang, Jason E McDermott, Jian-Ying Zhou, Vladislav A Petyuk, Li Chen, Debjit Ray, et al. Integrated proteogenomic characterization of human high-grade serous ovarian cancer. Cell, 166(3):755–765, 2016.

[20] Yuichi Abe, Maiko Nagano, Takahisa Kuga, Asa Tada, Junko Isoyama, Jun Adachi, and Takeshi Tomonaga. Deep phospho-and phosphotyrosine proteomics identified active kinases and phosphorylation networks in colorectal cancer cell lines resistant to cetuximab. Scientific reports, 7(1):1–12, 2017.

[21] Danica Wiredja. Phosphoproteomic Characterization of Systems-Wide Differential Signaling Induced by Small Molecule PP2A Activation. PhD thesis, Case Western Reserve University, 2018.

[22] Eric B Dammer, Andrew K Lee, Duc M Duong, Marla Gearing, James J Lah, Allan I Levey, and Nicholas T Seyfried. Quantitative phosphoproteomics of alzheimer’s disease reveals cross-talk between kinases and small heat shock proteins. Proteomics, 15(2-3):508–519, 2015.

[23] Cheng Chiang, Aleksander Tworak, Brian M Kevany, Bo Xu, Janice Mayne, Zhibin Ning, Daniel Figeys, and Krzysztof Palczewski. Quantitative phosphoproteomics reveals involvement of multiple signaling pathways in early phagocytosis by the retinal pigmented epithelium. Journal of Biological Chemistry, 292(48):19826–19839, 2017.

[24] Andrew Chatr-Aryamontri, Rose Oughtred, Lorrie Boucher, Jennifer Rust, Christie Chang, Nadine K Kolas, Lara O’Donnell, Sara Oster, Chandra Theesfeld, Adnane Sellam, et al. The biogrid interaction database: 2017 update. Nucleic acids research, 45(D1):D369–D379, 2017.

[25] Arthur Liberzon, Aravind Subramanian, Reid Pinchback, Helga Thorvaldsdottir, Pablo Tamayo, and Jill P Mesirov. Molecular signatures database (msigdb) 3.0. Bioinformatics, 27(12):1739–1740, 2011.

[26] Gerard Manning, David B Whyte, Ricardo Martinez, Tony Hunter, and Sucha Sudarsanam. The protein kinase complement of the human genome. Science, 298(5600):1912–1934, 2002.

[27] Aditya Grover and Jure Leskovec. node2vec: Scalable feature learning for networks. In Proceedings of the 22nd ACM SIGKDD international conference on Knowledge discovery and data mining, pages 855–864, 2016.

[28] Shaosheng Cao, Wei Lu, and Qiongkai Xu. Deep neural networks for learning graph representations. In Proceedings of the AAAI Conference on Artificial Intelligence, volume 30, 2016.

[29] Bryan Perozzi, Rami Al-Rfou, and Steven Skiena. Deepwalk: Online learning of social representations. In Proceedings of the 20th ACM SIGKDD international conference on Knowledge discovery and data mining, pages 701–710, 2014.

[30] Anton Tsitsulin, Davide Mottin, Panagiotis Karras, and Emmanuel Muller. Verse: Versatile graph embeddings from similarity measures. In Proceedings of the 2018 world wide web conference, pages 539–548, 2018.

[31] Martin Lee Miller, Lars Juhl Jensen, Francesca Diella, Claus Jørgensen, Michele Tinti, Lei Li, Marilyn Hsiung, Sirlester A Parker, Jennifer Bordeaux, Thomas Sicheritz-Ponten, et al. Linear motif atlas for phosphorylation-dependent signaling. Science signaling, 1(35):ra2–ra2, 2008.

[32] Vít Nováček, Gavin McGauran, David Matallanas, Adrian Vallejo Blanco, Piero Conca, Emir Munoz, Luca Costabello, Kamalesh Kanakaraj, Zeeshan Nawaz, Brian Walsh, et al. Accurate prediction of kinase-substrate networks using knowledge graphs. PLoS computational biology, 16(12):e1007578, 2020.

