## Supplementary material for "Utilization of Landscape of Kinases and Phosphosites To Predict Kinase-Substrate Association": Supp Fig 1

### Supplementary Materials

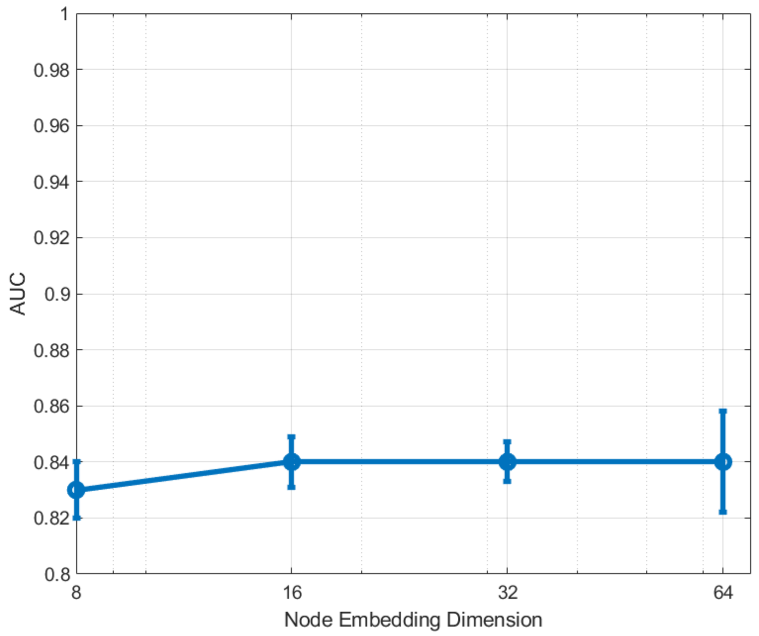


*Figure S1. Effect of number of dimensions of node embedding on prediction performance.*
